# Drug resistance-associated mutations in *Plasmodium* UBP-1 disrupt ubiquitin hydrolysis

**DOI:** 10.1101/2022.09.15.508122

**Authors:** Cameron Smith, Ryan Henrici, Maryia Karpiyevich, Megan R. Ansbro, Johanna Hoshizaki, Gerbrand J. van der Heden van Noort, David B. Ascher, Colin J. Sutherland, Marcus C.S. Lee, Katerina Artavanis-Tsakonas

## Abstract

Deubiquitinating enzymes function to cleave ubiquitin moieties from modified proteins, serving to maintain the pool of free ubiquitin in the cell while simultaneously impacting the fate and function of a target protein. Like all eukaryotes, *Plasmodium* parasites rely on the dynamic addition and removal of ubiquitin for their own growth and survival. While humans possess around 100 DUBs, *Plasmodium* contains ∼20 putative ubiquitin hydrolases, many of which bear little to no resemblance to those of other organisms. In this study, we characterize *Pf*UBP-1, a large ubiquitin hydrolase unique to *Plasmodium spp* that has been linked to endocytosis and drug resistance. We demonstrate its ubiquitin activity, linkage specificity and assess the repercussions of point mutations associated with drug resistance on catalytic activity and parasite fitness.

## INTRODUCTION

*Plasmodium*, like all other eukaryotes, deploy post-translational protein modifications as a means of maintaining homeostasis while driving the progression of its life-cycle. Components of the ubiquitin pathway mediate certain critical cellular processes, including protein degradation and trafficking, and are essential to parasite survival (1). The temporal and functional characterisation of ubiquitin pathway enzymes during *Plasmodium* development may uncover novel drug targets with the potential to provide new and urgently needed treatments for malaria.

Ubiquitin is attached to protein substrates via an enzymatic cascade consisting of an ATP-dependent E1 activating enzyme, an E2 conjugating enzyme and an E3 ligase. This modification is dynamic and can be reversed through the action of ubiquitin hydrolases (DUBs). *Plasmodium falciparum* expresses around 20 putative DUBs, annotated by homology with other eukaryotic enzymes, but the complexity and function of the malarial ubiquitin system remains mostly uncharacterized. UBP-1, a putative *Plasmodium* ubiquitin hydrolase, was first highlighted in *P. chabaudi* and linked to drug resistance (2). By sequencing *P. chabaudi* clones evolved *in vivo* in response to artesunate and chloroquine treatment, two separate valine to phenylalanine point mutations were identified in resistant parasites that both occurred within the C-terminal region of UBP-1. Two subsequent studies extended this link to *P. berghei* (3) and *P. falciparum* (4) through transgenic introduction of the corresponding point mutations into wild type lines, demonstrating that modified parasites had increased resistance to artemisinin, and in the case of *P. berghei* to chloroquine as well. Moreover, disruption of UBP-1 protein localisation through knock-sideways technology in lab-adapted *P. falciparum* also confers artemisinin resistance (5). Although the valine to phenylalanine mutations have not been identified in field isolates, other naturally occurring replacements within the *Pf*UBP-1 sequence, such as R3138H, have been correlated with artemisinin resistance (6).

The UBP-1 gene locus is located on chromosome 1 and consists of 3 exons encoding a large protein of 416 kDa. Although other *Plasmodium* species all appear to contain a UBP-1 homologue, higher eukaryotes lack homologues, barring low-identity hits to the catalytic domain alone. Transcriptomic analysis shows that it is predominantly transcribed in merozoites and rings (7), suggestive of a role in these early stages of infection. Indeed, UBP-1 appears to regulate the process of endocytosis, critical for the growth and maturation of the later blood stages. It localizes to the parasite cytostome, the site of initiation for uptake of host cell cytosol, and in parasites where UBP-1 has been perturbed, food vacuoles appear significantly smaller (5). The parasiticidal activity of artemisinin is dependent on efficient uptake and transfer of hemoglobin to the food vacuole during early ring stages of development. Once there, artemisinin is transformed to its active form by interacting with heme (8).Thus, impaired endocytosis would result in lower uptake of hemoglobin and link UBP-1 function to resistance. Together, these findings all point to a critical role for this protein in mediating drug susceptibility.

As the V3275F and V3306F mutations related to *in vitro* drug resistance occur near or within the UCH domain of UBP-1, we sought to explore its putative function as a ubiquitin hydrolase and to assess the effects of these mutations on catalytic function. Large scale mutagenesis identified *Pf*UBP-1 as being essential for parasite viability (1). However, disruption of the entire gene does not pinpoint the specific function or region of this protein that is required: is it the DUB activity or is it something related to the large upstream portion of *Pf*UBP-1? In this study, we aimed to characterize the ubiquitin hydrolase activity of *Pf*UBP-1, to assess the repercussions of these point mutations on catalytic activity and parasite fitness, and to gain insight into the function of this essential protein. We validate *Pf*UBP-1 as a true deubiquitinating enzyme with wide specificity for a variety of ubiquitin linkages. We further demonstrate that although V3275F and V3306F dramatically reduce catalytic activity, they leave parasite fitness unaffected. Our findings raise interesting questions about the putative function of UBP-1 and its role in mediating both endocytosis and artemisinin resistance.

## RESULTS

### *Pf*UBP-1 V to F mutations affect DUB activity but not fitness

*Plasmodium falciparum (Pf)*UBP-1 is a large, 3753 amino acid protein **(Figure 1A)** with a predicted UCH C19 peptidase domain at its C-terminus and no other recognizable motifs. Although overall sequence identity between UBP-1 orthologs of different *Plasmodium* species is relatively low, the UCH domain, and in particular the valine residues associated with drug resistance, are conserved **(Figure 1B)**. The modeled structure of the predicted C19 peptidase domain aligned closely with experimental structures of ubiquitin carboxyl-terminal hydrolases (sequence identities up to 33%, rmsd < 1 Å across residues 3181-3489), consistent with its proposed function, however the upstream region was predicted to be largely disordered and low-confidence.

**Figure 1.**
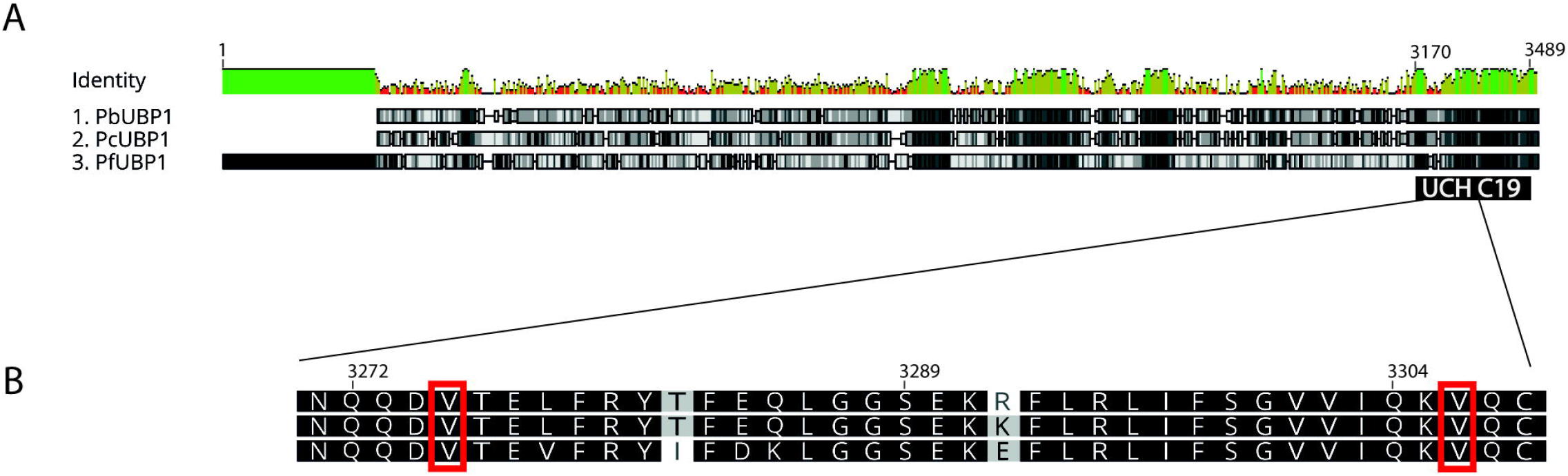
*Plasmodium spp*. UBP-1 alignment. (A) Alignment of the full sequence of UBP-1 from *Plasmodium berghei* (Pb), *Plasmodium chabaudi* (Pc) and *Plasmodium falciparum* (Pf) showing high (green) and low (red) identity regions and conserved amino acids in grayscale. (B) Detail of alignment of the UCH domain portion containing the two valine residues mutated in this study.

Considering both valine residues that are mutated in the drug-resistant parasites lie within the C19 peptidase domain, we hypothesized that introducing a valine at either position would result in protein misfolding and/or impair catalytic activity. Both V3275 and V3306 are buried and proximal to the active site cysteine, and their mutation to phenylalanine would result in a large change in overall residue volume within the tightly packed protein core. Modeling the mutations suggests that a change in either position would result in significant steric clashes (**Figure 2A-B)**, altering the local protein structure and potentially disrupting the active site architecture via changes to molecular packing and flexibility of the surrounding region. This was consistent with predictions that both mutations would lead to a significant decrease in protein stability (∼Kcal/mol), by SDM2, mCSM-Stability, DUET and DynaMut. Normal mode analysis indicated that both mutations would lead to altered protein dynamics and flexibility, which can be visualized as changes in the vibrational energy **(Figure 2C-D)**.

**Figure 2.**
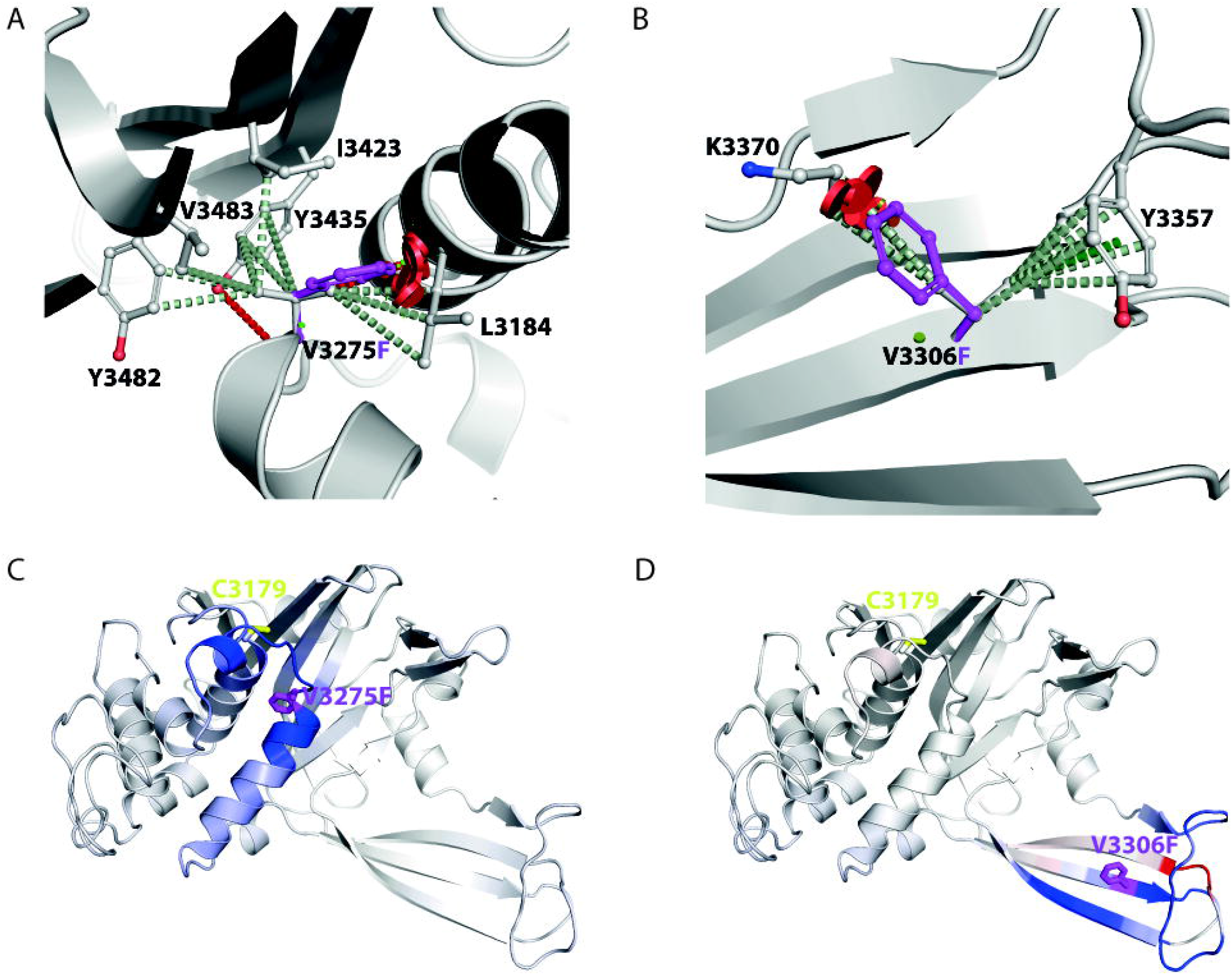
Molecular depiction of the effects on the *Pf*UBP-1 UCH domain caused by mutations V3275F and V3306F. The molecular interactions made by V3275 (A) and V3306F (B) were calculated and shown by Arpeggio as dashed lines. Hydrogen bonds are shown as red dashed lines, hydrobophic as light green and pi and deep green. Steric clashes caused by the mutations to phenylalanine are shown as red disks. The change in Vibrational Entropy Energy, calculated by DynaMut, from the wild-type to V3275F (C) and V3306F (D), where residues are coloured from blue, representing a rigidification of the structure, to red, a gain in flexibility.

Interestingly, the propagation of these changes were predicted to alter the dynamics near the active site cysteine.

As V3275F and V3306F transgenic parasites have been shown to be viable (4), we used these lines in a head-to-head competition assay to determine if our hypothesized impairment in DUB activity would result in more subtle differences in growth rates. V3275F, V3306F and the parental 3D7 line were each co-cultured at a 1:1 ratio with a fluorescent reference line. Over the course of three weeks and at regular intervals, mixtures were stained with Mitotracker Deep Red and the ratio of GFP positive to negative parasites was detected by flow cytometry to measure relative growth. Results showed equivalent rates of growth for both mutant lines as compared with the parental 3D7 **(Figure 3)** and, notably, no measurable reduction in fitness for either UCH-domain mutant.

**Figure 3.**
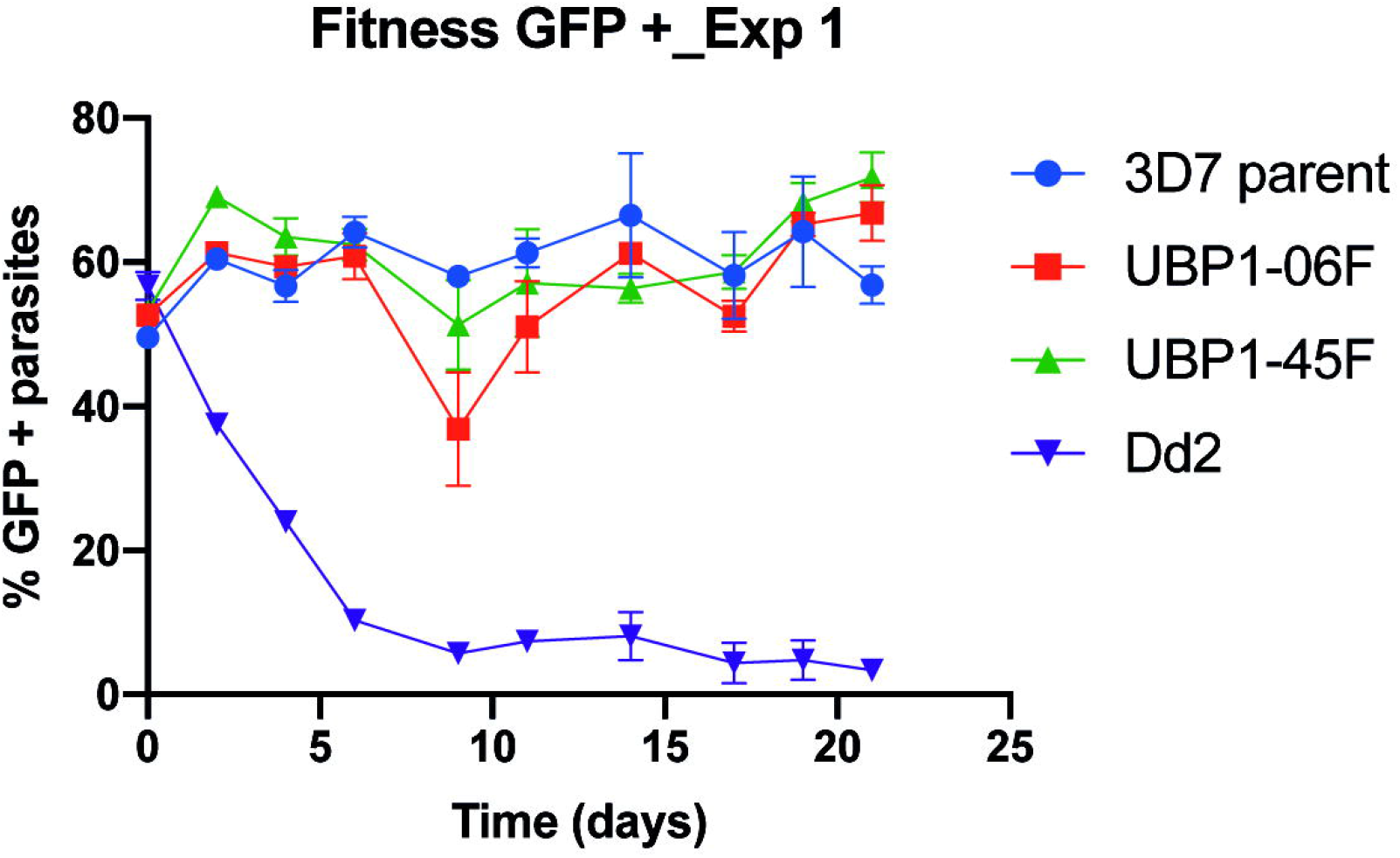
Fitness of UBP1 mutant parasites. UBP1 mutant lines V3275F and V3306F or wild-type 3D7 were individually competed against a fluorescent reporter line Dd2-EGFP. Parasites were mixed at a 1:1 ratio and the relative proportions of non-fluorescent test versus fluorescent reference line were measured by flow cytometry at regular intervals, with total parasites detected by staining with Mitotracker Deep Red. Shown is the average of three independent cultures, with error bars representing standard deviation.

### *Pf*UBP-1 contains a functional UCH domain

In light of these results, we questioned whether the DUB domain is indeed functional. To directly demonstrate deubiquitinating activity of UBP-1, we generated a truncation mutant of the UCH domain and used a bacterial expression system to produce recombinant protein. The truncation was designed to encompass as little extra sequence as possible outside the UCH domain while preserving alpha helices and structural integrity. Obtaining a measurable amount of protein was not trivial, as the entirety of UBP-1 is too large for bacterial expression and many truncations failed to express altogether. Several different combinations of truncations and bacterial strains were tested as well as an extensive array of expression, lysis, purification and buffer conditions. Ultimately, we were able to express a fragment containing the last 548 amino acids of UBP-1 with a molecular weight of 64.5 KDa (UBPtrunc) and through site-directed mutagenesis introduced the V3275F and V3306F mutations. We also mutated the active site Cys3179 to alanine to assess whether proteolytic function is dependent on the canonical Cys-His-Asp triad of the UCH domain.

Ubiquitin hydrolysis of UBPtrunc was first tested using a Ub-AMC fluorescence intensity assay which relies on real-time measurement of amido-methyl-coumarin (AMC) fluorophore release from the Ub reagent by the enzyme. In this assay, enzymes perform multiple catalytic cycles until all the Ub-AMC substrate is depleted, providing information on activity kinetics and allowing direct comparisons between mutants. The wild type construct generated a robust activity curve which flattened once substrate was depleted, whereas the mutant proteins failed to demonstrate any appreciable activity above baseline and behaved similarly to the C3179A negative control **(Figure 4A)**.

**Figure 4.**
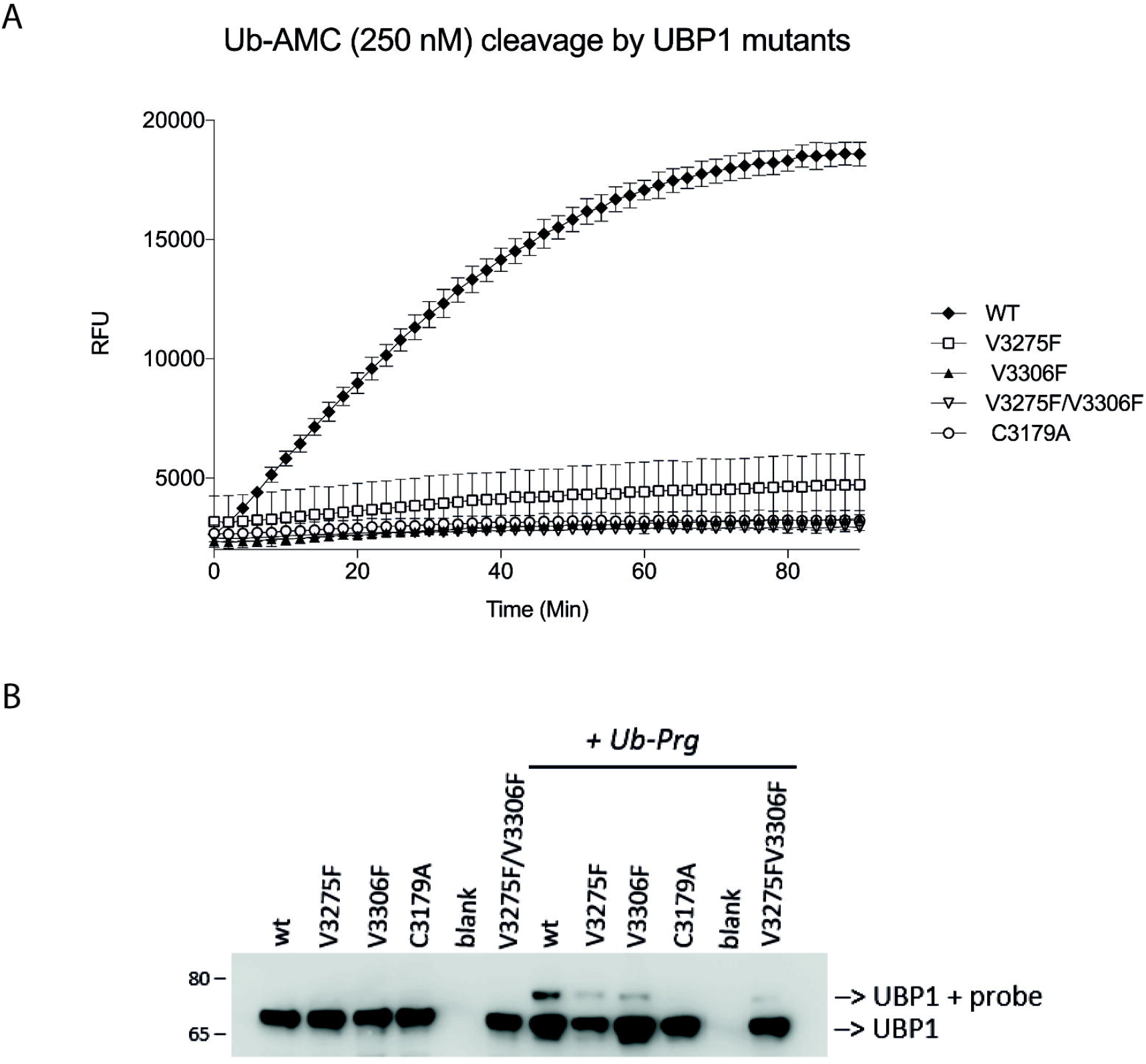
DUB activity of *Pf*UBP-1 wild type and V3275F and V3306F mutant proteins. The *Pf*UBP-1 C-terminal truncation (2952-3499) was expressed in wild type, V3275F, V3306F, C3179A and V3275F/V3306F forms with a 6xHIS tag. (A) Proteins were reacted with Ub-AMC substrate over the course of 1.5 hours (x-axis), with release of fluorescence (measured in relative fluorescence units, y-axis) indicative of activity. Reactions were done in triplicate. (B) Proteins were also reacted with a Ub-Prg probe. A shift up in electrophoretic mobility detected by anti-HIS immunoblot represents binding of the probe and activity.

To further investigate the mode of action of UBPtrunc and the effect of the mutations on the catalytic mechanism we reacted wild type and mutant proteins with HA-Ub-Prg, a ubiquitin derivatized probe designed to covalently bind the highly nucleophilic active site cysteine of deubiquitinating enzymes (18). An active ubiquitin hydrolase would be expected to form a covalent and non-cleavable complex with the probe and thus migrate at a higher molecular weight in SDS-PAGE analysis, as indeed is observed when reacting wt PfUBPtrunc with the probe (Figure 4, lane 7). The C3179A mutation rendered the UCH completely inactive as no complex formation was observed (lane 10). Both V3275F and V3306F mutants showed impaired complex formation by gel-shift assay and the double mutant showed an even more pronounced effect on complex formation, although in all V-to-F mutants some minor residual activity did remain **(Figure 4B)**. Of note, this type of assay results in the active site cysteine being covalently coupled to the Ub-based probe and hence each UBP-1 enzyme can only react once with the suicide probe.

Together, these results demonstrate that UBP-1 does contain a functional UCH domain but also that the V to F mutations do impair proteolytic activity as predicted by our models. A closer look at these predicted structures in **Figure 2** suggests two ways these mutations may affect the UBP-1 interaction with Ub. In the case of V3275F, the additional steric bulk of the phenylalanine within the small binding groove effectively narrows the access to the enzyme’s active site which may partially block Ub binding. On the other hand, V3306 seems to form part of a Ub-binding S1 pocket interface. Thus, the Val to Phe mutation at this position may disrupt the interaction between Ub and UBP-1 and thus the orientation necessary to allow catalysis.

### *Pf*UBP-1 has wide specificity for Ub linkages

Although mammalian UCH-DUBs are known to prefer cleaving Ub-from small peptide fragments rather than cleaving polyUb (or diUb) chains we wanted to assess whether this holds true for the *Pf*UBPtrunc (19–21). DUB activity of UBPtrunc was assessed by testing its ability to hydrolyse a panel of di-Ub linkage variants **(Figure 5A)**(22). These variants were made using the mammalian ubiquitin sequence which differs in one amino acid (D16E) with *Plasmodium* ubiquitin, however we have previously shown that this single substitution does not affect recognition by *Plasmodium* enzymes (23, 24). UBPtrunc was incubated with each of the seven isopeptide linked diUb variants as well as the linear M1 linked diUb followed by SDS-PAGE analysis and Coomassie Blue staining to visualize UBPtrunc mediated proteolysis of the dimeric Ub into monomeric Ub. Apart from confirming the activity of the UBPtrunc towards polyUb chains the results are also indicative of specificity for a particular subset of linkage types. The results showed that although *Pf*UBPtrunc cannot process linear ubiquitin chains (M1) and at most only marginally proteolyses K27 and K29 diUb, it is indeed a bona-fide deubiquitinating enzyme with a wide acceptance of isopeptide linkages, able to cleave K6, K11, K33, K48 and K63-linked di-Ub efficiently **(Figure 5B)**. Confirming that diUb proteolytic function is also dependent on the canonical Cys-His-Asp triad, the C3179A mutant showed complete loss of function for all diUb isotypes.

**Figure 5.**
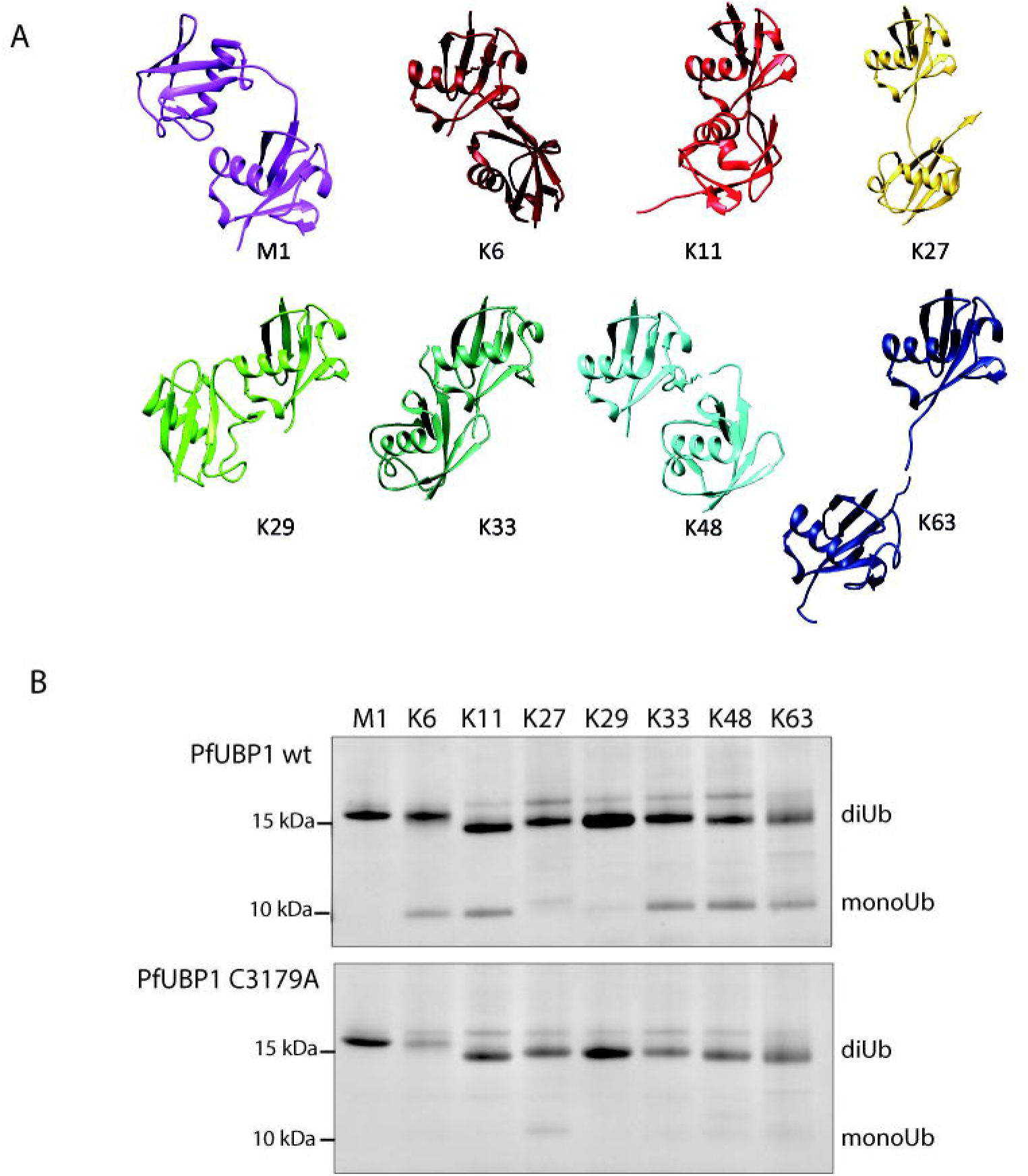
*Pf*UBP-1 linkage specificity. (A) The panel of di-Ub linkage isotypes used to assess specificity. Positions of attachment are depicted by each numbered lysine. (B) A truncation of the C-terminal end of the enzyme containing the UCH catalytic domain was expressed and mixed with each di-Ub individually. Activity was assessed by the release of monomeric Ub.

## DISCUSSION

In this study we demonstrate the DUB activity of *Pf*UBP-1 and its ability to hydrolyse a wide range of ubiquitin linkages. We also show that point mutations associated with drug resistance markedly interfere with enzyme activity *in vitro* and, in molecular modeling *in silico*, are predicted to disrupt the stability of the catalytic domain. These results would support DUB activity as being critical to UBP-1 function and to parasite homeostasis but, as the mutant *P. falciparum* lines displayed no measurable growth defect, that ablation of the DUB activity is not fatal for blood-stage growth. Thus the essential role of UBP-1, which prevents viability of knockout parasites, may require the upstream sequence, within or without a fully functional catalytic site. We are currently generating systematic truncations and plan to explore the minimal UBP-1 domain required for parasite viability in blood stages.

Our *a priori* assumption was that the DUB activity is what renders *Pf*UBP-1 essential for parasite growth. The lack of growth defect in parasite lines harboring either of the *Pf*UBP-1 mutants V3275F and V3306F was therefore unexpected. It has been established in *P. berghei* that UBP-1 is an essential gene for parasite survival, yet beyond the UCH C19 peptidase motif at its C-terminus, the upstream portion of its sequence contains no other recognisable domains that could explain these observations. Perhaps the V3275F and V3306F mutants retain low level DUB activity that is in fact sufficient to fulfill its essential functions, thus allowing the parasite to grow normally. Moreover, the presence of the entirety of the PfUBP-1 protein may somehow correct for subtle conformational changes making the point mutations less detrimental. The fact that both Val to Phe mutants retained some residual activity to the HA-Ub-Prg probe combined with our modeling results, would suggest that these mutations may serve to disrupt UBP-1 interaction with the Ub target, rather than ablating proteolytic efficiency. Generating a true catalytically dead mutant *in vivo* by converting C3179 to Ala would clarify this point, however all our attempts to do so were unsuccessful. The same CRISPR construct and guide that successfully installed both Val to Phe mutations failed to insert the C3179A in three separate experiments. Future studies could introduce this substitution using conditional induction in order to seek conclusive evidence of any link between failure to grow and catalytic loss.

We and others have linked *Pf*UBP-1-mediated reduced susceptibility to artemisinin with endocytosis in the early ring-stage trophozoite ((4, 5, 25). UBP-1 has been localized to the cytostome by microscopy and mislocalization of the protein results in disrupted delivery of host hemoglobin-rich cell cytosol to the food vacuole (5). In turn, reduced hemoglobin uptake has been linked to reduced drug susceptibility, with defective trafficking and protein degradation thought to mechanistically underlie artemisinin resistance (26). Our results demonstrate that *Pf*UBP-1 Val to Phe mutations adjacent to the catalytic site may ablate DUB function, and our previous results suggest parasites expressing these mutations are viable but artemisinin resistant. Taken together, these data suggest that the *Pf*UBP-1 DUB activity may play a central role in host cytosol uptake and delivery to the food vacuole in the early trophozoite, but over the full 48h intra-erythrocytic cycle, sufficient uptake occurs compensating for any *in vitro* fitness loss early in development. However, as these mutations have not been encountered in naturally-occurring *P. falciparum*, we cannot rule out a fitness deficit *in vivo*, and this may be due to additional roles for UBP-1 in liver, sexual or mosquito stages of the life cycle. DUB involvement in endocytosis has been described in higher eukaryotes, although these processes are notably distinct from the host cytosol ingestion of *Plasmodium* and, further, UBP-1 bears no resemblance to AMSH and UBPY, the mammalian enzymes involved (27). Of note, the Ub linkage preference profile of UBP-1 seems identical to that of UBPY (USP8) and more closely resembles that of other USP-type DUBs (manuscript submitted). Even though UBP-1 is predicted to have a UCH domain, it is possible that it interacts with ubiquitin in a manner more reminiscent of USPs (28). Further structural studies are needed to understand how it folds and its exact mode of binding to Ub.

Despite the evolutionary conservation of the ubiquitin-proteasome system across eukaryotes, few predicted *Plasmodium* DUBs appear to be direct orthologs of characterized enzymes in other organisms (24, 29). *Plasmodium’s* unusual biology undoubtedly requires the action of specialized proteins capable of supporting its unique growth and development. Similarly to UBP-1, most *Plasmodium* ubiquitin pathway enzymes are apparently essential (1). Defining their behavior, both temporally and functionally, will undoubtedly uncover a number of novel targets for antimalarial development and increase our understanding of cellular homeostasis in *Plasmodium* parasites.

## EXPERIMENTAL PROCEDURES

### Cloning and mutagenesis

The C-terminal region of PfUBP-1 (PF3D7_0104300) spanning amino acids 2952 - 3499 and containing the catalytic C19 peptidase domain was amplified from *P. falciparum* 3D7 cDNA (forward primer 5’-TATGGATCCCATATGATGAAAAATGTAAAAC-3’ and reverse primer 5’-TCTAAAGCTTTTAAAAGTACAAATCTGGAG-3’) and cloned into a SUMO-modified pet28a vector (kind gift from Dr. Owain Bryant) to yield a protein with a 6xHis and SUMO tag on the N-terminus. Point mutations C3179A, V3275F, and V3306F were introduced using the Quick-change II Site-Directed Mutagenesis Kit (Agilent, UK).

### Protein expression and purification

SoluBL21 *E*.*coli* (Genlantis, CA, USA) were used for expression of wild type and mutant *Pf*UBP-1 protein variants. Bacterial cultures were grown at 37°C, and protein expression was induced with 0.16 mM IPTG at OD_600_ of 0.6 for 16-20 hours at 18°C. Protein was purified under native conditions according to the QIAexpressionist (Qiagen) protocol. Bacterial pellets were resuspended in lysis buffer (50mM NaH_2_PO_4_, 300 mM NaCl, 10mM imidazole, 1 mM DTT, 5U/ml benzonase, 2 µg/ml aprotinin, pH8) and passed through a cell disruptor at 30 kpsi. The lysates were centrifuged at 40000g for 1 hour at 4°C. The supernatants were mixed with CoNTA beads (Generon) and incubated for 2 hours at 4°C on a rotary shaker. After the incubation, the beads were centrifuged, the flow-through was removed, and the beads were washed 6 times in wash buffer (50mM NaH_2_PO_4_, 300 mM NaCl, 10mM imidazole, pH8). As the attempts to elute *Pf*UBP-1 from the beads yielded inactive protein, all subsequent reactions were performed with *Pf*UBP-1 on beads. As such, concentrations were calculated by eluting bound protein from a defined volume of beads and quantifying by BCA (Pierce). Normalised volumes of beads bound to defined amounts of protein were then used in each assay.

### Di-Ubiquitin synthesis

All isopeptide linked di-Ubiquitin chains were generated using total chemical protein synthesis of monoUb mutants in analogy to the described procedure by El Oualid et all. (PMID: 22213387). In short, a Ub mutant equipped with a _γ_-thiolysine on the prospective lysine was reacted with a Ub-thioester in denaturing buffer (8 M Guanidium-HCl (Gdn.HCl)/100 mM phosphate buffer at pH 7.6 supplemented with 100 mM tris(2-carboxy-ethyl)phosphine (TCEP) and 100 mM 4-Mercaptophenylacetic acid (MPAA)). The crude di-Ubiquitin was first purified using RP-HPLC purification and subsequently desulphurised using 2,2’-Azobis[2-(2-imidazolin-2-yl)propane] Dihydrochloride (VA044) and reduced glutathione in 8 M Gdn.HCl/100 mM phosphate buffer with 100 mM TCEP at pH 7.0. Purification using RP-HPLC purification and S75 16/600 Sephadex size exclusion column (GE healthcare) chromatography using 20 mM TRIS, 150 mM NaCl at pH 7.6 resulted in the natively iso-peptide linked di-Ub that was concentrated by spin filtration and snap frozen and stored at -80 C until further use. M1-linked di-Ub was recombinantly expressed following the procedure earlier reported by Rohaim et all. (PMID: 22281738). All di-Ub are based on the human Ub sequence, which deviates at position 16 from the parasite sequence (human: E, plasmodium falciparum: D). This however has been shown to not impact recognition by plasmodium DUBs.

### Di-Ubiquitin cleavage assay

PfUBP-1 wild type or catalytic inactive PfUBP-1 C3179A mutant were incubated with 2 µg di-Ubiquitin in reaction buffer (50 mM NaH_2_PO_4_, 300 mM NaCl, 1 mM DTT) for 1 hour at 37°C. The reactions were terminated by adding 4X reducing sample buffer (Boston Biochem) supplemented with 100 mM DTT and loaded on gel for SDS-PAGE analysis. Visualization of the protein bands was performed using Coomassie Blue staining.

### Activity-based probe assay

PfUBP-1 wild type and mutant proteins were incubated with 10 µg of ubiquitin propargylamide (Ub-Prg) probe in reaction buffer (50 mM NaH_2_PO_4_, 300 mM NaCl, 1 mM DTT) for 1 hour at 37°C. The reactions were terminated by adding 4X reducing sample buffer (Boston Biochem) supplemented with 100 mM DTT.

### Amido-methyl-coumarin (AMC) assay

AMC assays were performed as described previously (Karpiyevich et al 2019 Plos Pathog). Briefly, the reactions were performed in AMC buffer (50 mM Tris (pH 7.4), 150 mM NaCl, 2 mM EDTA, 2 mM DTT, 1 mg/ml BSA). The reactions were initiated by the addition of 250 nM ubiquitin-7-amido-4-methylcoumarin (Ub-AMC, Boston Biochem) substrate to PfUBP-1 wt or catalytic inactive PfUBP-1 C3179A mutant. The fluorescence was measured on a BMG FLUOstar Omega plate reader continuously for 1 – 3 hours at room temperature using an excitation wavelength of 355 nm and an emission wavelength of 460 nm.

### Parasite growth

*P. falciparum* 3D7 parasites were grown in RPMI-1640 with 50 mg/l hypoxanthine, 0.25 % sodium bicarbonate, 0.5 % Albumax II, 25 mM HEPES (Life Technologies), supplemented with 1X concentration (2mM) GlutaMAX (Gibco) and 25 µg/ml gentamicin (Melford Laboratories). Human O+ RBCs were used for propagation of *P. falciparum* 3D7 parasites at 4% hematocrit.

### Competition assays

Growth of mutant parasites were directly assessed against a Dd2-EGFP reference line as previously described (9). Briefly, equal numbers of unmarked wild type (3D7), V3275F, and V3306F parasites were mixed with Dd2-EGFP and assessed for growth over time using flow cytometry. Mixtures were stained with Mitotracker Deep Red to detect all parasites, and ratios of non-fluorescent test line to fluorescent reference line were measured to calculate growth relative to the reference line.

### Immunoblot

Protein levels were normalised using BCA (Pierce) and separated by SDS-PAGE before semi-dry transfer to PVDF at constant 140mA for 1 hour. Membranes were blocked in 5% milk and probed with 3F10-HRP anti-HA antibody (Roche) followed by imaging with ECL.

### Structural modeling

A molecular model of the C-terminus of *Pf*UBP-1 (2952-3499) was generated by AlphaFold2 (10). The molecular consequences of the V3275F and V3306F were analyzed for their effects on the protein structure (11). The effects of the mutations on the stability of *Pf*UBP-1 were predicted using mCSM-Stability (12), DUET (13) and SDM (14). The effects of the mutations on protein stability, dynamics and flexibility were assessed using DynaMut (15) and DynaMut2 (16). The molecular interactions made by the wild type and mutant residues were calculated and visualized using Arpeggio (17).

## Acknowledgements

This work was supported by a Wellcome Trust Career Development Award WT085054MA and a BBSRC Project Grant BB/R001642/1 both held by KAT. CS was supported by a Wellcome Trust Doctoral Training Program studentship. CJS is supported by the UK Medical Research Council. DBA is supported by the National Health and Medical Research Council (NHMRC) of Australia (GNT1174405). ML was supported by Wellcome [Grant number 206194/Z/17/Z]. GJvdHvN acknowledges funding by NWO (VENI 722.01.002 & VIDI 192.011). For the purpose of open access, the author has applied a CC BY public copyright license to any Author Accepted Manuscript version arising from this submission.

